# Straining the root on and off triggers local calcium signaling

**DOI:** 10.1101/2023.05.11.540308

**Authors:** Vassanti Audemar, Yannick Guerringue, Joni Frederick, Isaty Melogno, Pauline Vinet, Avin Babataheri, Valérie Legué, Sébastien Thomine, Jean-Marie Frachisse

## Abstract

Throughout their life, plant root are submitted to mechanical stresses due to pressure exerted by the soil. So far, few studies addressed root cell deformation and calcium signaling elicited by soil compression. In this study, we designed a microchip inspired by pneumatic microvalve concept in order to deliver a lateral pressure to the root of a plant expressing the RGECO1-mTurquoise calcium reporter. Lateral pressure applied on the root induced a moderate elastic deformation of root cortical cells and elicited a multicomponent calcium signal at the onset of the pressure pulse, followed by a second one at the release of the pressure. This indicates that straining rather than stressing of tissues is relevant to trigger the calcium signal. The calcium elevation was restricted to the tissue under pressure and did not propagate. Additionally, the calcium signals exhibited a remarkable attenuation upon repetitive stimulations.

**Highlights:**

- A microvalve concept mimicking lateral soil pressure was developed.
- Non-damaging lateral compression of the root induces an elastic deformation of cortical cells.
- A multicomponent calcium signal is elicited at the onset of a pressure pulse and upon release of the pressure.
- Straining rather than stressing of tissues is relevant to trigger the calcium signal.
- The calcium signal is localized at the tissue under pressure and does not propagate.
- Calcium signals exhibit a remarkable attenuation upon repetitive stimulations.

## Introduction

Plants are anchored to the ground by their roots. They need to sense environmental cues to adapt to external conditions. Among these cues are external forces like gravity, soil compression, wind or touch by animals. In contrast to aerial organs, roots experience high mechanical stresses due to pressures exerted by the soil. In order to penetrate the soil and to overcome physical obstacles, the root generates an axial force. At the same time, during its progression, lateral confinement along the radial axis increases, generating lateral forces [1], [2]. Soils scientists often consider “soil structure” as the spatial arrangement of the different components and properties of soil [3]. A typical volume of surface soil includes about 50% solids, mostly soil particles (45%), and organic matter (generally < 5%) and about 50% pore space [1]. Therefore, during their growth, roots progress in a heterogeneous network crossing empty cavities and substrates of various stiffness. The thrust force (or pushing force) exerted by the growing part of the root has to overcome the soil resistance as well as the lateral friction with the soil. The friction involved in the balance of forces is the one acting on the flanks of the root along the elongation and meristematic zones [1]. A local lateral confinement around the radial axis is a scenario that the root can encounter. Such a stress occurs for example upon radial growth of the root squeezed between two hard fixed soil particles. To our knowledge, the characteristics of physical and biological responses locally elicited by such a compressive, non-wounding stimulation, have so far not been investigated.

Calcium is one of the most important ions for signal transduction. Free cytosolic calcium concentration increases in response to many signals. The duration, amplitude, frequency and spatial distribution of the calcium elevation is controlled by calcium channels, transporters and pumps localized at the cell membranes [4]. The spatio-temporal pattern of cytosolic calcium elevation was shown to encode information allowing specific responses to diverse cues that involve cytosolic calcium as a second messenger [5], [6]. Notably, it has been shown that calcium is involved in signal transduction of touch [7]. Rise and propagation of a calcium signal in the case of Venus flytrap induces the closing of the leaf [8]. *Arabidopsis thaliana* also displays a calcium signal after local stimulation of a root cell with the tip of a micropipette [7]. Macromolecules involved in the control of the wall integrity embedded in the membrane or in the cell wall could be recruited for transduction of a mechanical stress into biological responses including calcium variations [9]. For example, FERRONIA (FER), a transmembrane protein was shown to be involved in root mechanoperception [10]. Moreover the *fer* mutant shows an alteration of the calcium signal elicited by touching or bending the root [10]. The plasma membrane is also subjected to mechanical stress due to tensile or compressive forces and variation of osmotic pressure [11]. Calcium permeable mechanosensitive channels at the plasma membrane are good candidates to mediate cytosolic calcium elevations in response to membrane deformation induced by touch or cell compression. Thus far, five families of mechanosensitive channels MSL, Piezo, OSCA, MCA and TPK have been molecularly identified and electrophysiologically characterized in Arabidopsis [12]. An additional mechanosensitive channel, non-molecularly identified, called RMA (Rapid Mechanically Activated) from the plasma membrane of Arabidopsis was characterized [13]. All these mechanosensitive channels except for MSL are calcium permeable but also permeable for other divalent and monovalent cations. With rapid activation and inactivation, Osca, Piezo and RMA share common kinetics properties [14].

Here we address the following questions: Could a compression mimicking the lateral confinement generated by the soil pressure deform the root? What are the specific properties of the calcium signal elicited by such strain? We developed a microfluidic device enabling us to apply a controlled mechanical lateral stress on the root to address these questions. The microfluidic device allows imaging of plant roots with a microscope for long durations with a high spatio-temporal resolution. We used confocal microscopy to image and quantify cell deformation and Epifluorescence microscopy combined with a fluorescent cytosolic calcium reporter to characterize the calcium signal induced by lateral strain on *Arabidopsis thaliana* roots.

## Materials and Methods

### Microfluidic device manufacturing

The polydimethylsiloxane (PDMS) devices were made using standard dry film and soft lithographic procedures based on the method by Dangla *et al*. (2013) [15]. To produce molds for the root growth channels, two layers of Eternal Laminar E8020 dry photoresist film of thickness 49± 2 μm were successively deposited on a glass slide by lamination at 100 °C to reach a desired channel height of ∼100 µm. The film was UV exposed through a photomask designed using CleWin5, to produce straight channels with height, width and length of ∼90 μm, 600 μm and 2 cm, respectively.

The channels were replicated from the master molds in degassed PDMS with a 1 to 10 ratio of curing agent to bulk material (SYLGARD 184 elastomer and curing agent, Dow Corning), cured at 70 °C for 2 hours to obtain the device pieces. PDMS blocks serving as root growth channels were replicated with a strictly controlled height, so that once bonded to a glass coverslip, the top part of the channel leaves a PDMS membrane with a thickness of 250 or 460 µm which serves as a deformable push-down valve. PDMS blocks serving as pressure channels sitting above the plant growth channels were made using the same procedure, and with a thickness of 4 mm +/- 1 mm. A spin coater (Model WS-650MZ-23NPPB, Laurell) was used to obtain a thin layer of PDMS to coat coverslips. Several devices with different membrane thicknesses were produced by varying the time or rotational speed of the spin coating process. Membrane thicknesses were measured by profilometry (ProFilm3D, Filmetrix) (Supplementary Fig. 1). PDMS pieces were peeled off the molds, and pierced with 1mm holes to create liquid and gas inlets/outlets. Additional holes were punched at a 45° angle to serve as the entry path connecting the root with the root growth channels before sealing by plasma treatment (Harrick Plasma, Plasma Cleaner PDC-002-CE) together and to a glass coverslip covered with a thin (37 +/- 2 µm) PDMS layer.

### Plant material and growth conditions

Seeds of Arabidopsis thaliana (Col-0 ecotype) constitutively expressing RGECO1-mTurquoise under the UBQ10 promoter [16] were sterilized in ethanol 70% and SDS 0.05% for 5 min, rinsed with ethanol 96% for 5 min and dried at room temperature. Then, the seeds were sown on conical cylinders produced by cutting micropipette tips containing Hoagland medium (1.5 mM Ca(NO_2_)_2_, 0.28 mM KH_2_PO_4_, 0.75 mM MgSO_4_, 1.25 mM KNO_3_, 0.5 µM CuSO_4_, 1 µM ZnSO_4_, 5 µM MnSO_4_, 25 µM H_3_BO_3_, 0.1 µM Na_2_MoO_4_, 50 µM KCl, 3 mM MES, 10 µM Fe-HBED, pH 5.7) with 1% phyto-agar, they were inserted into the same medium filling Petri dishes [17]. After 3 days of stratification at 4°C in the dark, the seeds were incubated in 16h light/ 8h dark at 22°C during 3 days in a culture chamber. After the primary root reached the bottom of the cone, they were transferred from the Petri dish to the microfluidic device (Fig. 1b) and kept under the same temperature and light conditions. Root growth was conducted into the root channel filled with liquid Hoagland. During the growth, the device was tilted with an angle > 45° to allow the root to grow gravitropically. Root channels were connected by tubing to syringes filled with liquid Hoagland medium, and the channel medium was refreshed with a flow rate of 1 μL/min using a syringe pump.

**Figure 1:**
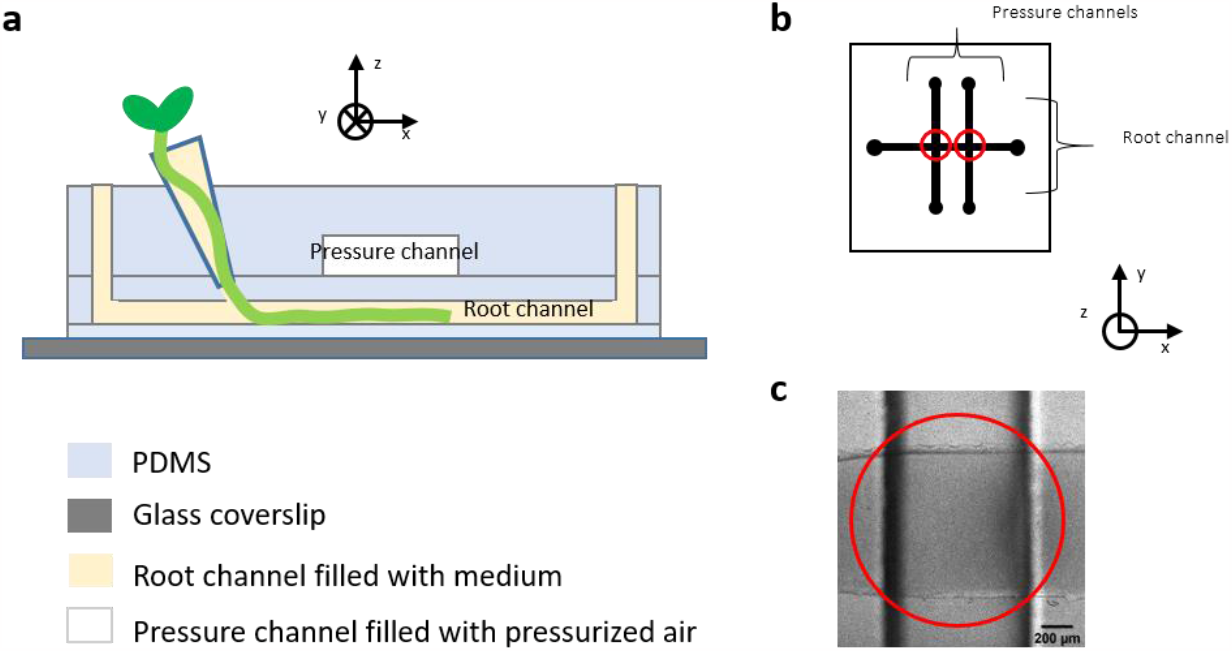
Microfluidic device equipped with a pneumatic valve. a) Schematic side view of the microfluidic device containing the plant root channel and pressure channel seprated by a flexible push-down PDMS valve. b) Schematic top view of the 2 mechanical push-down valves at the intersections of the root channel and the pressure channels. c) Top view of a mechanical push-down valve imaged in bright field (Leica DMI 6000, 5x objective). In 1.c and 1.d, valves areas have been circled in red.

### Acquisition protocol

When root growth had extended passed the deformable membrane portion of the PDMS device, i.e. 6 or 7 days after transfer to the incubation chamber, microscopy experiments were launched. The root channel was connected by tubing to syringes filled with liquid Hoagland medium and connected to a syringe pump (WPI, AL-1000) enabling control of the flow rate. The pressure channels were connected by tubing to a pressure box (Fluigent, MFCS-EX) that allows injection of an air flow at a fixed pressure into these channels. The PDMS device was secured in a 3D printed sample holder fitted with a Perspex lid and designed to fit in a standard 96-well plate sample holder.

Cross-sectional views of the microfluidic channels and cross sections of the roots were acquired with a Leica SP8 inverted microscope equipped with, a white light laser (470 to 670 nm), and two GaAsP Hybrid detectors (Hamamatsu). For cross sections of the channels, fluorescein solutions at 10 µM were imaged with a 10x PLAN APO dry objective (Leica) at λex = 488 nm and λem = 501-609 nm. For cross section of the roots, cell walls were labelled with propidium iodide (5 µg/mL) and imaged using a 20x PLAN APO multi-immersion objective (Leica), with λexc = 488 nm and λem = 551-651 nm. A Leica DMI 6000 inverted microscope equipped with an excitation lamp (PE-4000 LEDs, CoolLed), a quad band dichroic mirror (Chroma) and black and white camera (CoolSNAP HQ^2^ CCD, Photometrics) was used to image intracellular calcium. RGECO1-mTurquoise fluorescent lines were imaged using a 5x dry objective with λexc = 580 nm and λem = 600-700 nm for RGECO1 and λexc = 470 nm and λem = 490- 520 nm for mTurquoise.

### Image processing and data analysis

Image processing and analysis was performed using Matlab. Length variations of cells along Oy and Oz axis (Fig. 3) were measured on cross-sectional views of wild type roots stained with propidium iodide. Image analysis was conducted following these steps : the background was subtracted and a segmentation was performed to delimit cell boundaries. Maximal length of each cell in the horizontal and vertical dimension was measured before and during the pressure stimulation using bounding box (Supplementary Fig. 2). Calcium signal variation measurements were performed as follows : for each time point, the background was subtracted and a binary image was generated. The root axis was extracted and segments of 100 µm perpendicular and centered around this axis were distributed at 50 pixels intervals (Supplementary Fig. 2d). Mean values of the ratio between RGECO1 images and mTurquoise images were calculated for each segment along the root and reported in heat maps representing calcium signal variations along the root axis over time.

## Results

### Setting up a micromechanical system for delivering lateral pressure

In order to apply a controlled lateral compression to the root, we have developed a microfluidic device combining a rootchip [17] with a pressure system inspired by a micromechanical push-up valve [18]. The device was fabricated in PDMS which has been shown to be biocompatible with *Arabidopsis thaliana* [19]. Three layers of PDMS were sealed together, enabling the formation of channels: the pressure channels containing air sit on top of the root channels in which the root is growing, while the whole PDMS device is bound to a glass coverslip covered in a thin PDMS film to enable the visualization of the root with an inverted microscope (Fig. 1a).

The perpendicular layering of the root channel and pressure channels defines a square PDMS membrane (Fig. 1b) with an active area of 600 µm by 600 µm (Fig. 1c). The PDMS deformability and the specific dimensions, especially the thickness/side length aspect ratio, allow the membrane to deflect downward into the root channel when a sufficiently high pressure is injected into the pressure channel. In our device, two of these micromechanical push-down valves are distributed 2 mm apart over the root channel (Fig. 1b), and each pressure channel can be controlled individually.

We compared the deflection, **d**, of the push-down membrane of microfluidic devices with 2 different membrane thicknesses: 460 +/- 57 µm and 250 +/-31 µm (as annotated in figure 2a). The 250 µm PDMS membrane exhibits a greater deformability for the same pressure (shown in cross-sectional views of the root channels without root perfused with a solution of fluorescein at 10 µM with a constant flow rate of 8 µL/min; supplementary Fig. 3). A compromise had to be made for the PDMS membrane thickness to be thin enough to increase the deformability and enable a good transfer of pressure from the pressure channel to the root, but thick enough to avoid damage during the device manufacturing process. In further experiments, the PDMS membrane thickness was fixed at 250 μm.

**Figure 2:**
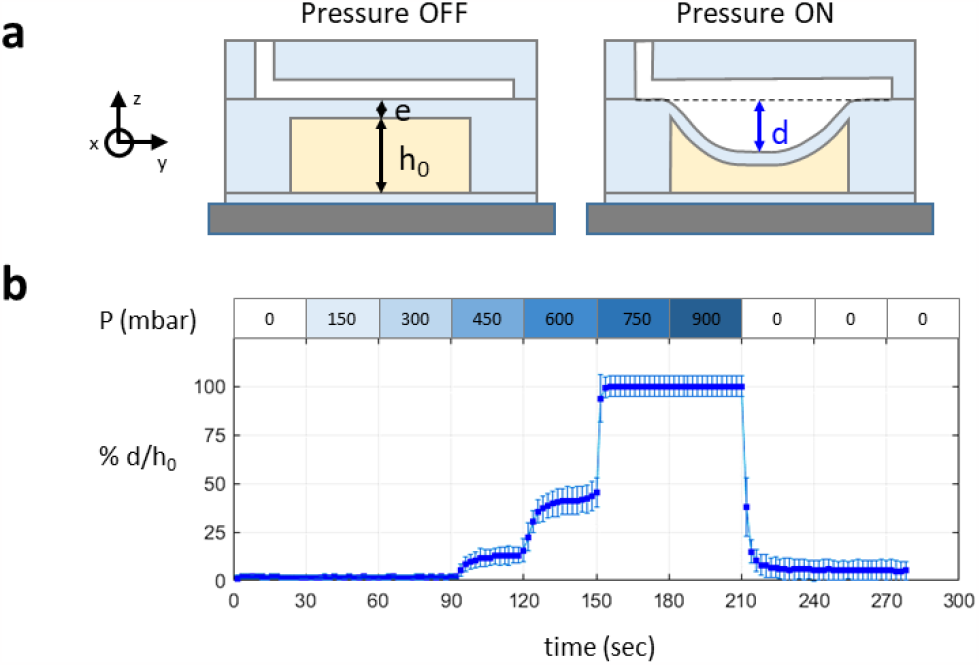
Relationship between membrane deformation and pressure in the air channel. a) Schematic cross section of the pneumatic valve with and without pressure: **e** is the thickness of the PDMS membrane separating the pressure channel (white) and the root channel (yellow), **h**_**0**_ is the height of the root channel before pressure and is equal to 90 μm, **d** is the maximum distance between the center of the valve membrane at rest and in its deformed state when pressure is applied. b) Variations of **d**, the maximal deformation of a 250 µm thick PDMS membrane with applied pressure. Measurements were repeated 3 times on 4 different valves. Squares correspond to the mean values of all measurements and error bars correspond to standard deviations.

**Figure 3:**
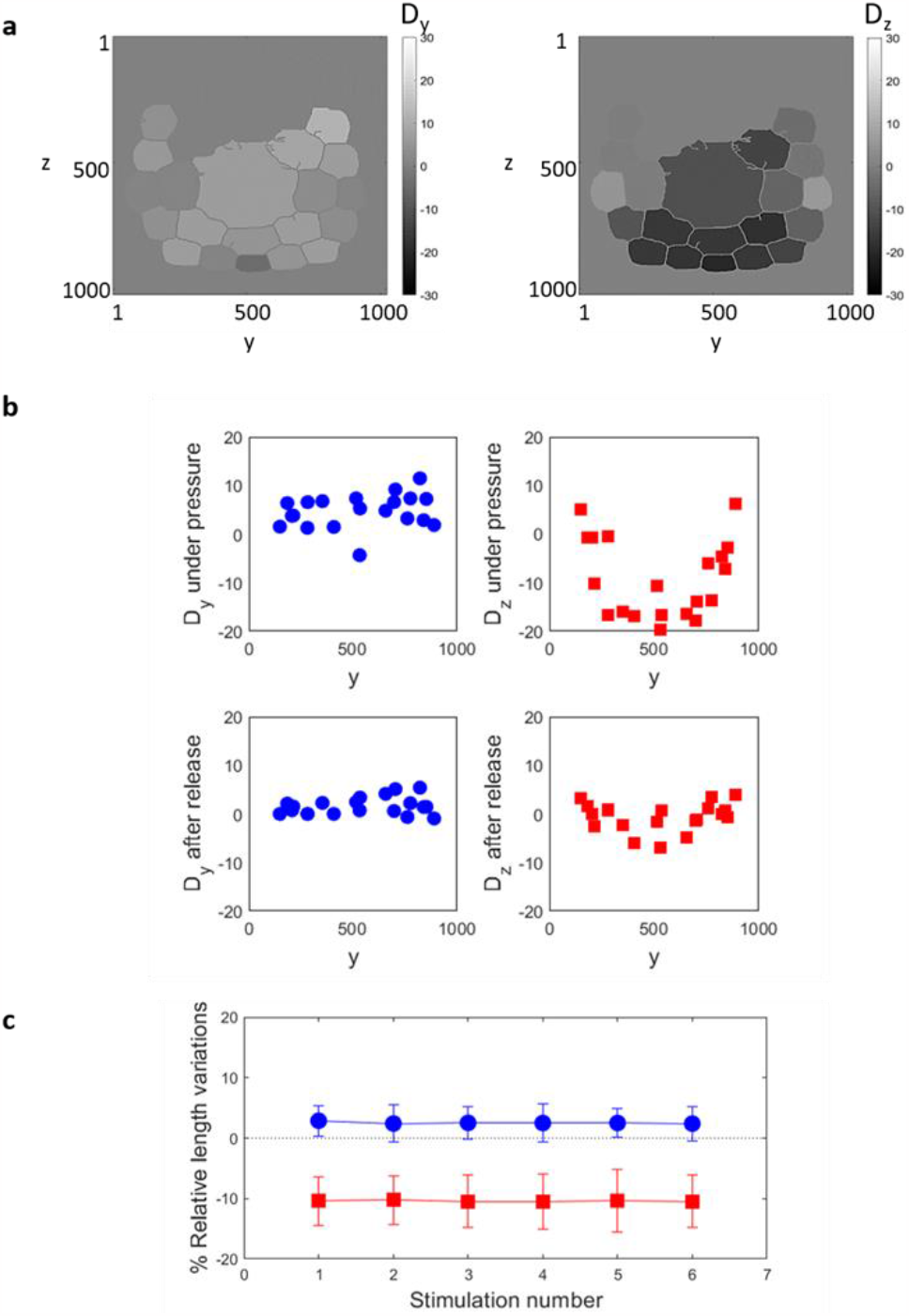
Transversal and longitudinal relative length variations D_y_ and D_z_: a) Representation of the segmented cells of one representative root under a pressure stimulation of 900 mbar. Each cell was colored with a grey level corresponding to the percentages D_y_ on the left and D_z_ on the right. b) Percentage D_y_ and D_z_ of one representative root during the first stimulation and after the release in function of the transversal dimension Oy. c) Percentages D_y_ and D_z_ averaged over all the segmented cells of a root stimulated 6 times every 5 min with a pressure of 900 mbar during 1 min. The plain symbols represent the mean value of 19 different roots and the error bars represent the standard deviation associated. For b) and c), the blue circles and red squares correspond to D_y_ and D_z_ respectively.

We measured the deflection of the 250 µm PDMS membrane as a function of the pressure increasing by steps of 150 mbar every 30 seconds. The percentage of the deflection length, **d**, normalized by the thickness of the channel **h**_**0**_ is presented in figure 2b. Deformation is triggered with pressure from 450 mPa and continuously increases with pressure, until the valve membrane reaches the bottom of the channel (d = h0) at 750 mPa. This system without root has a typical time to reach the equilibrium state of around 10s. Upon release of the pressure, the membrane returns to its approximate initial position. The number of stimulations on the same valve does not impact its ability to deform. The short standard deviation bars indicate that the system is reliable and does not experience damage with repeated stimulations.

### Lateral pressure induces quasi-elastic deformation of the root cells

We assessed the mechanical response to a pressure of 900 mbar applied through the deformable PDMS membrane (also called valve) on primary roots of 7 days old seedlings in the maturation zone (1-5 mm from the apex). Confocal images of root cross sections were used to measure the deformation of the root in the Oyz dimension. Cell walls were stained with propidium iodide to visualize cell shape. A segmentation analysis was performed to identify the outlines of cells (Supplementary figure 1a, Supplementary video 1). We measured the length of the cells, L_y_ and L_z_ projected on the Oy and Oz axes, transversal and longitudinal to the applied force respectively. D_y_ and D_z_ are the corresponding relative length variations defined by D_y_ = (L_y_ – L_y0_)/L_y0_ and D_z_ = (L_z_ – L_z0_)/L_z0_, with L_y0_ and L_z0_ the projected length on Oy and Oz axes before the application of the pressure. Cells located in the upper part of the image were out of the working distance of the objective and therefore could not be segmented. Cell walls located in the central cylinder were not stained by propidium iodide and therefore were not segmented, thus we consider the central cylinder as one object.

Under pressure, D_y_ and D_z_ show a heterogeneous repartition between the cells of the cross-section due to the geometry of the applied force and the connectivity between the cells (figure 3a), resulting in a complex tension field. For example, the cells located on the lateral sides experience a positive variation length D_z_, indicating an elongation instead of the compression observed among the other cells. Figure 3b shows the repartition of D_y_ and D_z_ along the transversal axis O_y_ for the root under pressure and after release. The distribution of D_y_ is roughly constant along the O_y_ direction whereas Dz experiences a symmetrical distribution with respect to the midline of the root in the Oz direction, with the largest relative length variations corresponding to the cells close to the midline (fig. 3a). After release of the pressure, the initial cell shapes were recovered with values of D_y_ and D_z_ close to 0 %, showing a quasi-reversible deformation process.

Six repeated stimulations every 5 minutes have been performed on roots with a pressure of 900 mbar during 60 seconds, on 19 roots in the maturation zone (1-5 mm from the apex). The average percentages of D_y_ and D_z_ over all the cells per root were calculated. No correlation was revealed between the average percentage of length variations and the position along the main axis. Under a stimulus of 900 mbar, D_z_ was around 10 % and D_y_ around 4%. After 6 stimulations, no significant differences in relative length variation was found as shown in figure 3c.

### Mechanical stimulus elicits a local elevation of cytosolic calcium concentration

We used the fluorescent ratiometric reporter RGECO1-mTurquoise [16] to monitor cytosolic calcium variations in roots. The reporter was constitutively expressed into the cytosol of plant cells, and we measured the ratio R between the RGECO1 fluorescent signal, sensitive to the calcium concentration, and the mTurquoise fluorescent signal, used as a control of the expression of the reporter. The normalized ratio R / R_0_, with R_0_ corresponding to the baseline value of R before any stimulation, was represented by heat maps in Fig. 4b, as a function of the time and the position along the main root axis for two types of stimulation at 900 mbar: a short stimulation of 30 s, or a long stimulation of 20 min. In either case, we observed a calcium signal rapidly increasing and progressively decreasing back to the basal level approximately 15 min after the beginning of the stimulation (Supplementary video 2). In the case of the long stimulation, a second increase of calcium signal is elicited by the release of pressure. On a spatial level, the calcium signal mainly stays between the boundaries of the valve and no clear propagation along the root axis was noticed. The normalized projected area A / A_0_ and the average value of the ratio 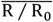 were calculated on the portion of the root located under the valve and are represented in Figure 4c with the pressure protocol associated. In either case, the variation of the projected area was around 5% and recovered its initial level after the release of pressure, indicating a small elastic deformation. The average value of the ratio 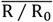 displays a multicomponent signal with two distinguishable phases: a rapid and sharp elevation with a maximum occurring at 10 ± 3.7 s after the beginning of the stimulation, followed by a smoother rise and decrease of the signal with a maximum a 100 ± 15.5 s, that ends up recovering its initial value within 15 minutes.

**Figure 4:**
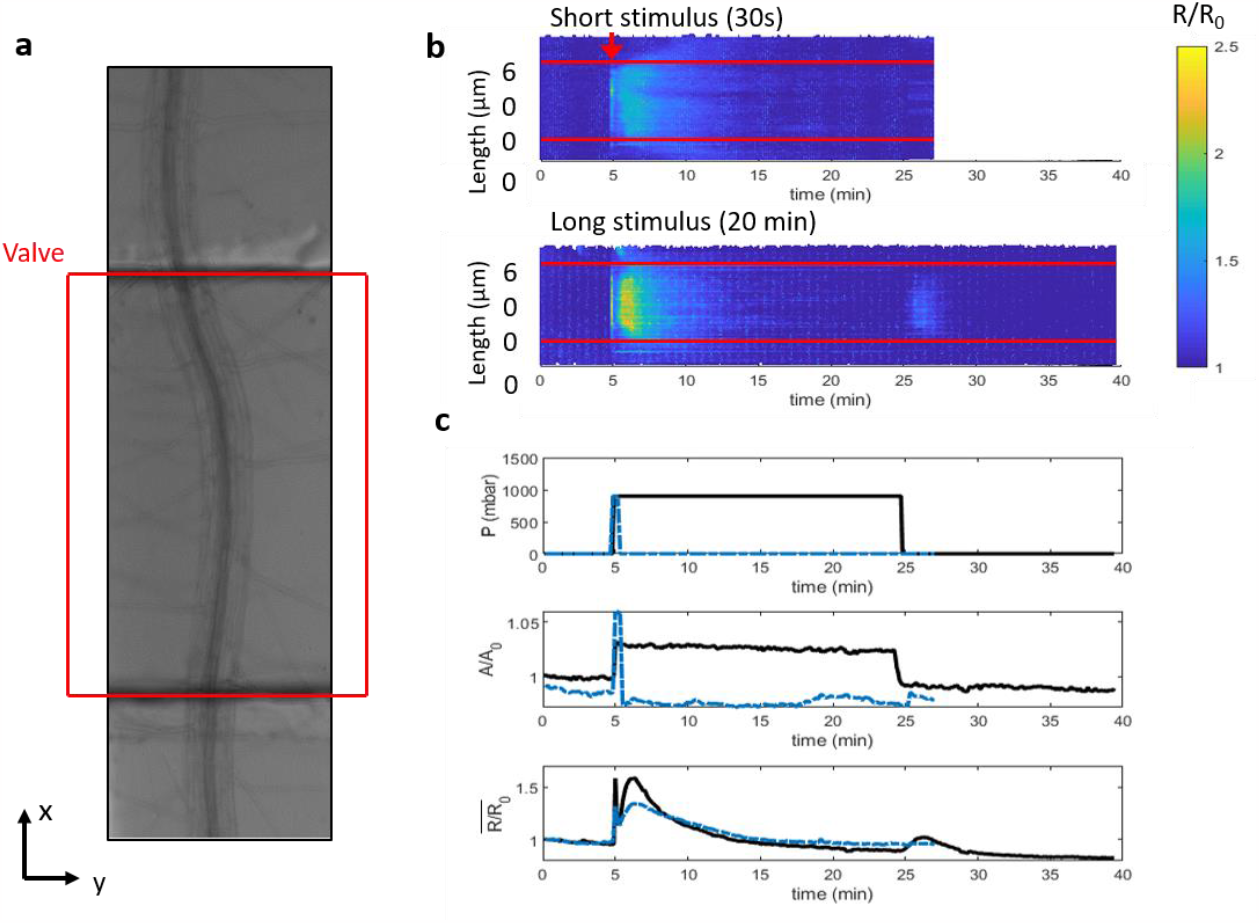
Calcium response, in time and space, to a local short (30 s) and long (20 min) applied pressure of 900 mbar: a) Bright field image of the root under the pneumatic valve. b) Variations of the normalized ratio R/R_0_ along the main root axis x during time, represented as heat maps for a 30 s stimulus (top image) and a 20 min stimulus (bottom image). The red lines correspond to the position of the pneumatic valve and the red arrows to the time of stimulation (activation and release). c) Time variations of the pressure stimulation (top), the normalized projected area of the root (middle) and the calcium signal ratio normalized to the value at time 0 (bottom) elicited by a pressure of 900 mbar during 30s (dotted blue line) and 20 min (black line). The graphs show one representative experiment out of 23 for short and 10 for long stimulation.

### The amplitude of the cytosolic calcium signal increases with increasing pressure intensity

In order to analyse the relationship between the pressure intensity and the amplitude of the cytosolic calcium signal, we subjected the root to mechanical stimulations of 30 s with an increasing pressure intensity using the pressure protocol presented in Figure 5a (top panel). The normalized area presented in Figure 5a (middle panel) show an increase corresponding to the increasing pressure. We can also observe the increase of the amplitude of the calcium signal as represented in the figure 5b. Note that the amplitude of the initial fast increase was not measured because the acquisition rate in this experiment was not sufficient to resolve it. However, repetition of the pressure stimulation at 750 mbar (for 2 plants) or further increase to 900 mbar (1 plant, Fig. 5) triggered a rise in cytosolic calcium with a lower amplitude. This indicates that the system is indeed sensitive to the intensity of pressure and suggests that it undergoes attenuation upon repetitive stimulations.

**Figure 5:**
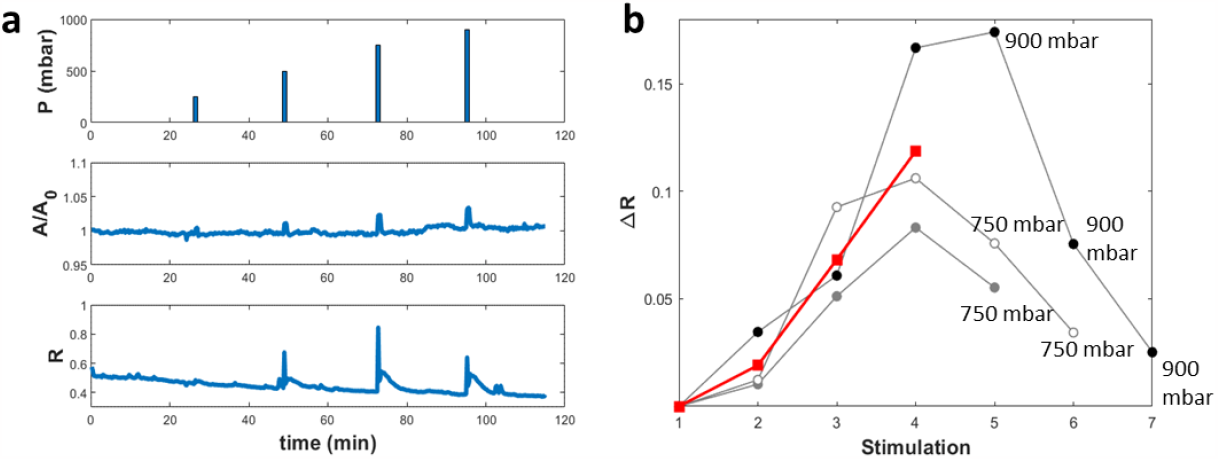
Successive stimulations of 30 s with an increasing pressure: a) pressure (top), normalized area (middle) and calcium signal ratio (bottom) for one representative plant. b) Amplitude of the calcium signal for each stimulation in three different roots represented in black, grey and white disks. Stimulations 2, 3 and 4 correspond to 250, 500, and 750 mbar, respectively. Stimulations 5 to 7: pressure as indicated on the graph. The red squares correspond to the mean value for the first 3 stimulations at 250, 500 and 750 mbar.

### Repeated stimulations lead to attenuation of the calcium signal

To test whether the decrease of the calcium signal upon increasing pressure stimulation (Fig. 5) is due to an intrinsic attenuation, we tested the effect of repetitive stimulations of the same amplitude. In Figure 6, we subjected roots to repetitive pressure stimulations of 30 s at 900 mbar every 5 min. The calcium signal in response to the first pressure pulse presented the highest intensity. The amplitude of the second peak was decreased by more than 50% compared to the first one, while the amplitude of following peaks subsequently decreased following each stimulation. These results indicate that the system displays attenuation upon repeated stimulation. To test the effect of the frequency of stimulation, and whether the system recovers from habituation after a longer delay, we imposed 30 second stimulations at different time intervals. In Figure 6C, the time interval between two pulses was increased to 20 min or 60 min. The amplitude of the calcium rise decreased irrespective of the delay between two stimulations. This indicates that the relevant factor for habituation is the number of stimulations rather than their frequency, and that the calcium dynamics system does not recover after 60 min.

**Figure 6:**
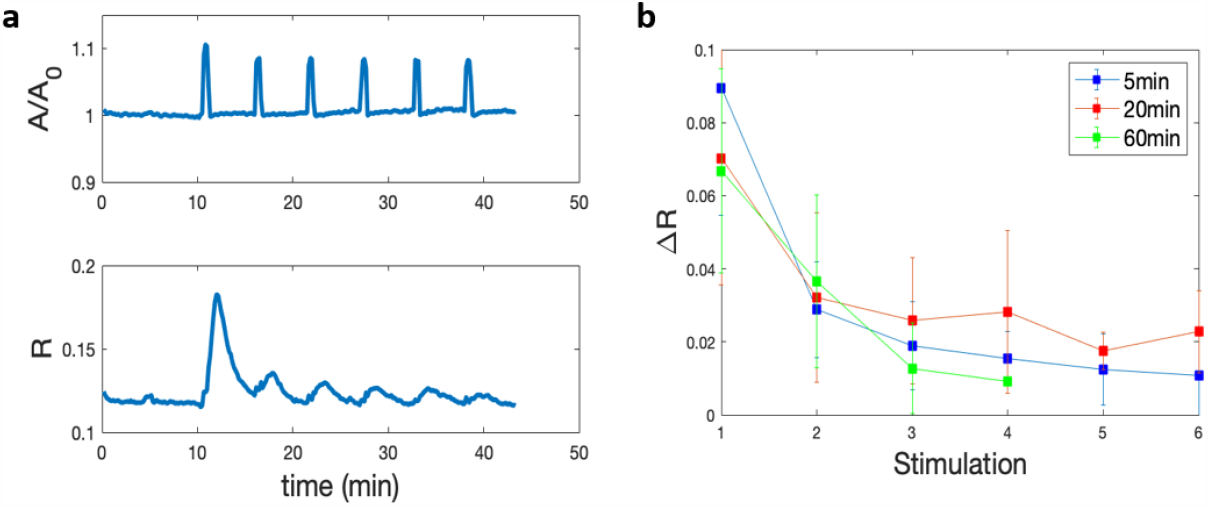
Repeated pressure stimulations lead to a decrease in the amplitude of the calcium signal. a) normalized area variations (top) and calcium signal ratio (bottom) in a representative experiment, b) mean value of maximum amplitude (+/- SD) of the calcium ratio peaks for each stimulation of 30 s at 900 mbar every 5 min (blue, n = 6), 20 min (red, n = 5) and 60 min (green n = 4).

### A progressive rise of pressure drives a different calcium signature

To apply a pressure stimulation that may correspond better to the stimulation experienced by a root growing between hard obstacles, we applied a progressive increase of pressure by steps of 150 mbar from 0 to 900 mbar every 60 seconds (Fig. 7). This stepwise increase was followed by a fast release of pressure. We repeated this protocol two times on the same portion of the root with a time interval of 20 min. In this configuration, the area deformation shows a stepwise increase and decrease after release of pressure corresponding exactly to the stepwise variations of pressure. A total recovery of deformation was observed after release of the pressure. The second stepwise stimulation gave rise to the same deformation. Compared to a one step increase, the calcium signal displayed a totally different shape with of slow integrated increase and no decomposition in two phases, and a lower amplitude. This shape could be due to an overlap between the rise and the decrease of the calcium signals triggered at each step. When the pressure was released from 900 mbar to 0 in a short time scale, a rapid increase was observed followed by a slow decrease to reach the baseline about 20 min later. Upon the second stepwise stimulation, the calcium increase appeared only at the highest pressure and its amplitude was reduced. This suggests that the calcium signal attenuation is associated with a decreased sensitivity to pressure.

**Figure 7:**
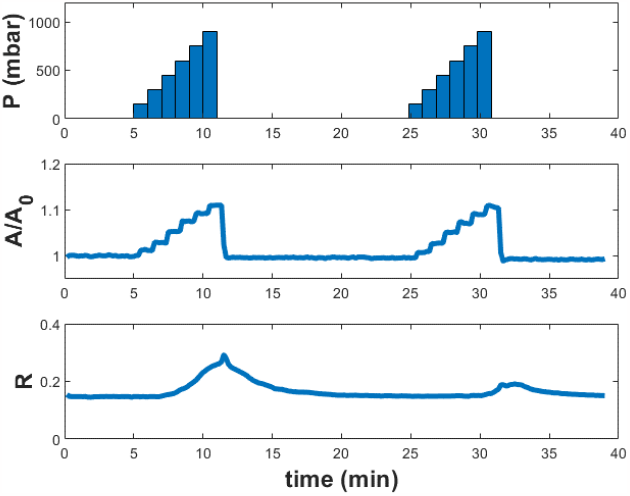
Calcium response to stepwise increases in lateral compression of the root. (top) Pressure increasing by steps of 150 mbar from 0 to 900 mbar every 60 seconds followed by release to 0 mbar. This stimulation was performed 2 times with a 20 min interval on a single root, (middle) Area variations with pressure stimulation over time, (bottom) calcium signal ratio over time.

## Discussion

We designed a microfluidic device allowing reproducible mechanical stimulations of roots. Our experimental design proved to be suitable for investigating root deformation together with calcium variation induced by a mechanical constraint. Precise description of the strain at tissue and cellular levels, generated by a local compression was achieved. The analysis of the cytosolic calcium concentration, in terms of kinetics, intensity and tissue location allowed us to characterize the Ca^2+^ signal and to link it with the local strain.

This system could be used to monitor a wide range of cellular parameters and events in response to gentle pressure stimulation. For example, the characterization of organelle shape and position upon mechanical stress would bring valuable information. Considering the variety of fluorescent probes now available and the ability to label proteins involved in mechanosensing, our system should allow addressing key questions in mechanotransduction such as, for example, microtubule reorganization and membrane tension variation.

### The calcium signature in response to mechanical stress

We observed a calcium signal composed of two components, a fast calcium increase peaking after a few seconds and a slower calcium response lasting a few minutes (Fig. 5 and 7), which are not propagated along the root. Monshausen et al. [7] also showed that both touching the surface of the Arabidopsis root or bending the root elicited a local calcium elevation. In both cases, the cytosolic calcium concentration rapidly increased and then returned to its initial concentration 10 minutes after bending and 60 seconds after touching. Mousavi et al. [20] monitored the calcium response to non-damaging mechanical indentation of the root cap of Arabidopsis. Likewise, increasing the amplitude of the indentations elicited a transient and localized Ca^2+^ signal in the columella and lateral root-cap cells. In addition, they showed that depleting the plant from the mechanosensitive channel PIEZO1 diminished the calcium transient [20]. When bending Arabidopsis root, Shih et al. [10] elicited a biphasic calcium response composed of a short peak followed by a longer wave. The longer wave was attributed to the activation of the Receptor-like Kinase FERONIA. A biphasic calcium response was also elicited by ATP in Arabidopsis roots. Matthus et al. [21] attributed the slow component to the activation of the plasma membrane receptor DORN1/P2K1, while the rapid component was attributed to the mechanical perturbation of the root through the experimental system. The calcium signature elicited by gentle mechanical stimulation is distinct from the signal induced by salt stress or wounding. Local treatment of the root with NaCl triggers Ca^2+^ waves that propagate through the plant at rates of up to ∼400 μm/s [22]. Calcium signaling induced by wounding the aerial parts or the root elicited a propagated wave of calcium delivering an information to remote organs [23].

### Compared to previous studies, our microfluidic set-up revealed new features of the calcium response

(i) a calcium elevation is observed upon increase but also upon release of the pressure, (ii) the intensity of the calcium response increases with the pressure applied and (iii) Successive pressure stimuli lead to attenuation of the calcium signal.

#### Attenuation

We have shown two important properties of the calcium signal, (1) after a rapid raise of calcium the signal slowly recovers its initial level after 10 minutes whether the pressure is sustained or not, and (2) the repetition of stimulations leads to a decrease in the amplitude of the calcium signal. Rapid increase in calcium, in response to various physical stimuli such as cold shock, osmotic shock or touch has been reported in plants [7], [24], [25]. These three stimuli, when sustained, the calcium concentration rapidly decreases within a few tens of seconds to a few minutes after the initial peak. In Arabidopsis and tobacco plantlets, the cold-induced increase in calcium is attributed to an influx of Ca^2+^ from the extracellular medium relayed by an intracellular store. Then, recovery would be due to ER and vacuolar uptake of calcium from the cytosol [24]. In Arabidopsis guard cells, performing successive depolarizing hyperosmotic KCl shocks, the authors showed that cytosolic Ca^2+^ concentration controls stomatal closure by two mechanisms, a short term ‘calcium-reactive’ closure and a long-term ‘calcium programmed’ steady-state closure [25]. Furthermore, similar to our results, an attenuation of the calcium signal is noticed upon repetition of the 5 minute KCl shocks every 10 minutes.

As part of an integrative approach, Martin et al. [26] have addressed the effect of wind stress on plant growth and gene activation by performing multiple stem bending on young poplars. They observed a decrease of the molecular response to subsequent bending as soon as a second bending was applied. They called this phenomenon desensitization and determined a refractory period of 7 days needed to recover a gene expression activation similar to that observed after a single bending.

The processes of amplification, attenuation and desensitization of the electrical signal have been the most investigated in the neural system in the context of the transmission of information. In neurons, theaction potential (AP) might fire up to a frequency of 500 Hz. In this system, the transmissionof signals via chemical synapses represents a very dynamic process. The synapse, playing a role of relay, is able in situations of prolonged stimulation to attenuate the signal [27]. The purpose of such an attenuation would be to process the information and to adapt to an excess of signal, such as mentioned for the auditory system [28]. The shape and the amplitude of the signal can be directly modified by the ion channels generating the AP. For example, in the case of a high stimulation frequency, some channels still being in their relative refractory period, APs will be modified in their shape and/or their frequency [29]. It is also exemplified by Cain et al. [30] who showed that the kinetic properties of several isoforms of T-type calcium channels are closely linked to their contribution to neuronal firing. Mutations of T-type calcium channels would be associated with certain pathophysiological disorders. The examples of attenuation mentioned above operate at different time scales, from milliseconds for APs in a nerve, to minutes in guard cells and roots of Arabidopsis, and up to days for poplar gene expression. Nevertheless, in all examples mentioned, attenuation leads to an adaptive response of the cell/organ/organism, and the primary actors of the generation of the signal are likely ion channels.

### Strain-stretch hypothesis, its physiological relevance

Calcium release is triggered at the onset of pressure and at the release of the pulse of pressure. Similarly, responses to “touch” and “letting go” have been reported for epidermis cells of Arabidopsis and tobacco [31]. In that case, distinct characteristics of the waves elicited by the compressive force and its release suggest different underlying mechanisms for the “touch” and “letting go». In our experiments, calcium release corresponds to the time when the maximum of strain variation in the root cells is observed. Indeed, tissue shape variation occurs when strain is applied and released. Although they differ in amplitude, probably due to attenuation, the calcium waves elicited by pressure and release share the same characteristics. This indicates that strain rather than stress triggers calcium signals underpinned by a common mechanism.

In biological materials, stress is not proportional to strain, therefore stress-sensing and strain-sensing mechanisms have different output [32]. For example, in Arabidopsis pavement cells, microtubules, which align in the direction of maximal mechanical stress, are postulated to play a role as a mechanosensor [33]. However, James et al. reported that more generally the stimulus for growth sensed by cells is the mechanical strain rather than the stress [34]. Furthermore, in agreement with our finding, Moulia et al. showed that the strain-sensing model is better suited than the stress-sensing model to explain the primary and secondary thigmomorphogenic growth-responses in trees [35].

Several recent studies highlight the role of calcium in long distance systemic signaling. Calcium signaling induced by wounding aerial part or root elicited a propagating wave of calcium delivering information to remote organs [23]. Here, with gentle local pressure the signal is restricted to the pressurized zone. Only strained tissues display a calcium signal. This signaling path probably indicates that cells have to react and adapt to the local deformation of the root. Thus, when a root is squeezed between hard objects such as stones, lateral tissues are likely pressure-stimulated, inducing a local calcium signal allowing the plant to adapt to this local soil constraint.

### What could be the molecular mechanisms underlying the calcium increase?

Although the calcium signature in response to mechanical cues displays common features, the molecular players remain elusive. Not only are the number of candidate receptors and channels possibly involved in Ca^2+^ response numerous, but also calcium sources are diverse. Indeed, many Ca^2+^ reservoirs are present in the cell, notably the vacuole, the endoplasmic reticulum (ER) and other organelles [36], [37]. The kinetic behavior of transient Ca^2+^ signals was modeled at the cell level and proposed to result from four components, two Ca^2+^ permeable channels located at the plasma- and endo-membrane, respectively, and two active Ca^2+^ efflux systems, a Plasma membrane -based Ca^2+^ ATPase pump and endomembrane-based Ca^2+^/H^+^ exchanger [4]. In our case, one can hypothesize that the short peak relies on the activation of mechanosensitive channels, which immediately activate upon membrane tension. The slower Ca^2+^ variation might recruit internal stores of calcium (ER, vacuole, …) governed by receptors involved in mechanosensing, such as FERONIA or DORN1.

At the cell membrane scale, Ca^2+^ permeable channels activated by a force applied in the plane of the membrane were recently identified and characterized. These channels, such as RMA-DEK dependent channels or those belonging to Osca and Piezo families, behave as transducers able to convert instantaneously a mechanical force into a Ca^2+^ flux [13], [20], [38]. One might hypothesize that a pressure locally exerted on the root induces tissue strain that leads to membrane stretching. In reaction to membrane stretching, calcium permeable mechanosensitive channels will be activated. Most of these channels (RMA, Osca, Piezo) present inactivation properties, meaning that a rise of pressure applied to the membrane activates the channel, but under sustained pressure the channel enters a non-conductive state called inactivated [14]. This inactivation might, at least in part, explain the rapid (1-2 min) decrease of the Ca^2+^ signal under a long pulse of pressure delivered by the valve. In our proposed scheme, the decrease of effective response to repetitive stimulation, that we call attenuation, would be provided by a mechanical modification of cellular structural elements. An increase in the cell stiffness would limit membrane stretching and then produce less activation of calcium permeable mechanosensitive channels. Whether the cytosolic Ca^2+^ elevation plays a role in this feedback loop remains to be investigated. Even though Ca^2+^ permeable force gated channels appear to be the best candidates to mediate the coupling between mechanical strain and cytosolic Ca^2+^ increase, other sensors of mechanical strains may also be involved, for example, the cytoskeleton itself, or sensors of cell wall integrity [9], [39], [40].

### What could be the adaptive outcome of local calcium signaling?

A root growing in the soil squeezed between two rocks, for example, will have to locally adapt its mechanical properties by strengthening its tissues. This could be achieved through strengthening the cell wall or remodeling the cytoskeleton. The non-propagated Ca^2+^ signal locally initiated by pressure stimulation (mimicked in the experiment represented in Figure 7) might be the event that initiates further transduction signaling cascades possibly involving pH variation, reactive oxygen species emission, and kinase activation, and further leading to developmental responses and to the root adaptation [5], [6]. In natural conditions, roots are also subjected to diurnal hydraulic pressure variations producing a periodic root diameter increase and decrease ([41]). This latter phenomenon has to be considered together with root progression in which a root squeezed in a bottleneck will be self-stimulated during growth. Root curvature additionally induces strains, and secondary roots preferentially emerge in the curved zones of the root [42]. This could be another adaptive response triggered by pressure stimulation of the root.

## Acknowledgements

Seeds expressing R-GECO were provided by Rainer Waadt and Melanie Krebs (Ruprecht-Karls-Universit#x00E4;t Heidelberg, Germany). This work has benefited from a French State grant (Saclay Plant Sciences, reference n° ANR-17-EUR-0007, EUR SPS-GSR) under a France 2030 program (reference n° ANR-11-IDEX-0003) through the DYNANO project. It has also benefited from Imagerie-Gif core facility supported by l’Agence Nationale de la Recherche (ANR-11-EQPX-0029/Morphoscope, ANR-10-INBS-04/FranceBioImaging ; ANR-11-IDEX-0003-02/ Saclay Plant Sciences). We gratefully acknowledge the financial support from the Région Ile de France through the DIM ELICIT program, for the Plantuidics grant. We thank David Bouchez from IJPB (Versailles, France) for fruitful discussions and Nicolas Valentin for printing adapters to set up microfluidic chips of the microscopes.

## Supplementary data

**S-Fig 1:**
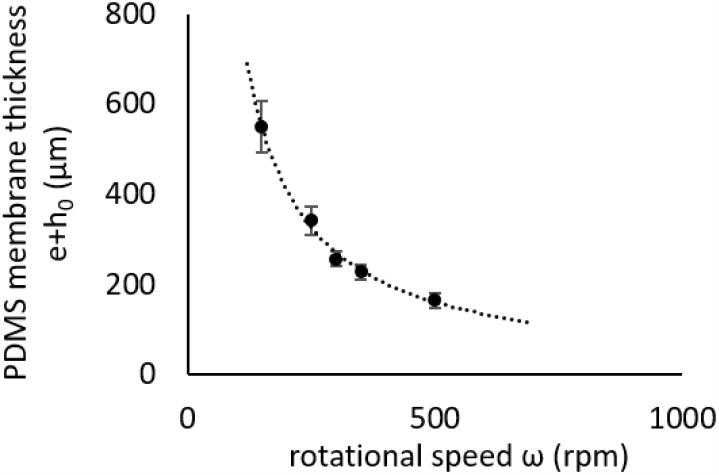
PDMS membrane thickness variations with ω, the rotational speed during the spin coating process. Circles correspond to experimental data and the dotted line corresponds to the power law fit given by the equation (e+h_0_) = 0.09ω^-1,02^. The fit of the curve with a power law is in agreement with Koschwanez et al. [1] and Zhang et al. [2].

**S-Fig 2:**
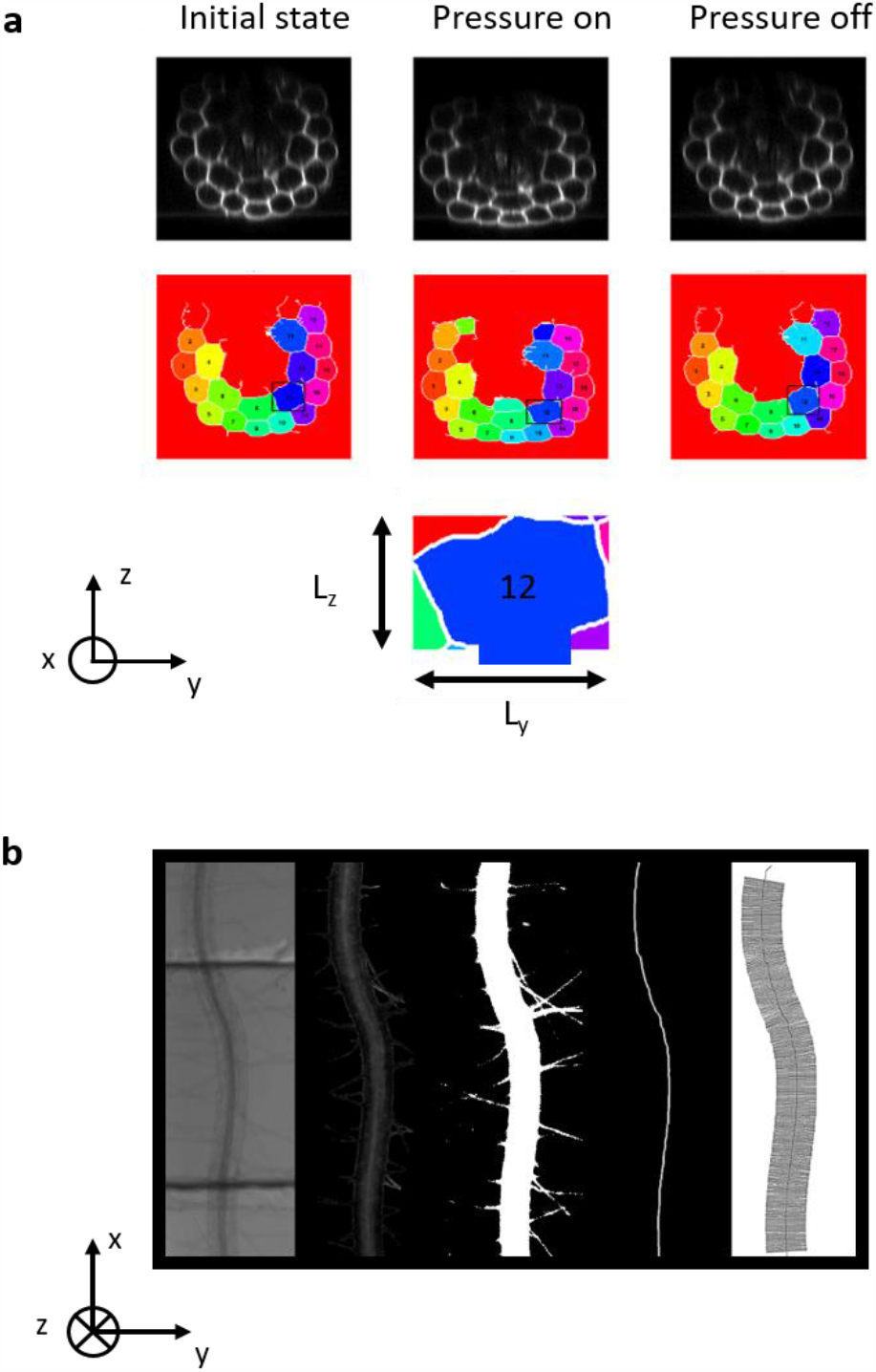
Image analysis procedure: a) Cross-sectional image of the root cells at rest, under a lateral pressure of 900 mbar and after release of the pressure; Top images (black and white), fluorescent images with cell walls colored with propidium iodide (5 µg/mL) and imaged with λexc = 488 nm and λem = 551-651 nm, Middle images (colored), segmented cells separated with white lines, colors have no other purpose than differentiating cells between each other. Cell number 12 is framed by a black bounding box and represented in the bottom line. The maximum height L_z_ and the maximum width L_y_ of the cell are indicated. b) Top view image of the root under the valve (from left to right) : bright field image, mTurquoise fluorescent image (λexc = 470 nm and λem = 490-520 nm), binarized image, and skeletonized axis of the root, segments (100 µm length) perpendicular to the root axis and distributed every 50 pixels.

**S-Fig 3:**
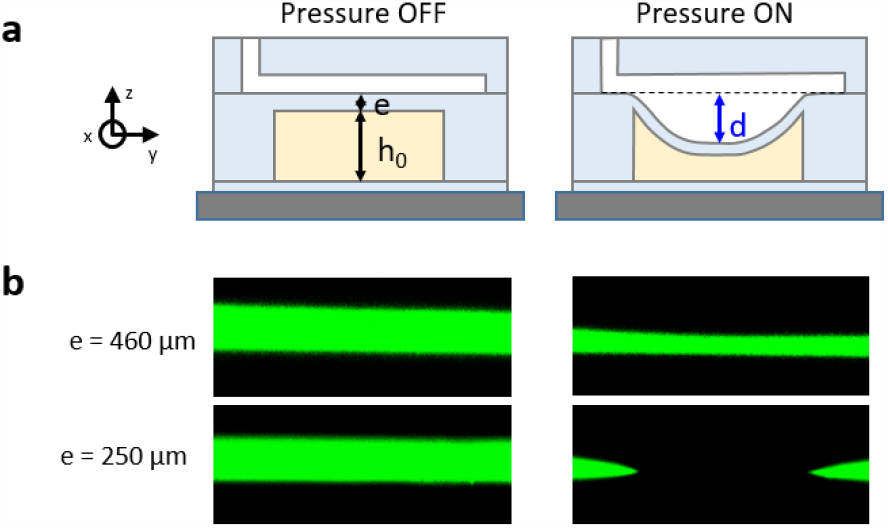
a) Schematic cross section of the PDMS device showing deformation of the pneumatic valve with and without pressure: e is the thickness of the PDMS membrane separating pressure channel (in white) and the root channel (in yellow), h0 is the thickness of the root channel without pressure and is equal to 90 μm, d is the maximum distance between the center of the valve membrane at rest and in its deformed state. b) Cross-sectional view of the microfluidics channels filled with a dilute solution of fluorescein without (left) and with (right) 750 mbar of pressure for two different PDMS membrane thicknesses: e = 460 μm (top image), e= 250 µm (bottom image). Deformability of the PDMS membrane decreases with its thickness.

## SUPPLEMENTARY VIDEO 1

Deformation of the root stimulated 6 times every 5 min with a pressure of 900 mbar during 1 min. Cross-sectional video of the root with cell walls colored with propidium iodide (5 µg/mL) and imaged with λexc = 488 nm and λem = 551-651 nm. The pressure is applied from the top and the playing speed is accelerated 50 times

## SUPPLEMENTARY VIDEO 2

Calcium response of the root to a local pressure of 900 mbar applied during 30 sec (short stimulation). Variations of the normalized ratio RGECO1-mTurquoise fluorescence (λexc and λem of RGECO1 and mTurquoise, see Material and Method).The dotted line correspond to the position of the pneumatic valve. The playing speed is accelerated 100 times

## SUPPLEMENTARY VIDEO 3

Calcium response of the root to a local pressure of 900 mbar applied during 20 min (long stimulation). Variations of the normalized ratio RGECO1-mTurquoise fluorescence (λexc and λem of RGECO1 and mTurquoise, see Material and Method).The dotted line correspond to the position of the pneumatic valve. The playing speed is accelerated 100 times

